# iRGD-liposomes enhance tumor delivery and therapeutic efficacy of antisense oligonucleotide drugs against primary prostate cancer and bone metastasis

**DOI:** 10.1101/2021.01.08.426005

**Authors:** Jibin Guan, Hong Guo, Tang Tang, Yihan Wang, Yushuang Wei, Punit Seth, Yingming Li, Scott M Dehm, Erkki Ruoslahti, Hong-Bo Pang

**Affiliations:** Department of Pharmaceutics, School of Pharmacy, University of Minnesota, Minneapolis, MN 55455, United States; Ionis Pharmaceuticals, Inc. Carlsbad, CA 92010, United States; Masonic Cancer Center, University of Minnesota, Minneapolis, MN 55455, United States; Departments of Laboratory Medicine and Pathology and Urology, University of Minnesota, Minneapolis, MN 55455, United States; Cancer Center, Sanford Burnham Prebys Medical Discovery Institute, La Jolla, CA 92037, United States; Cend Therapeutics, San Diego, CA 92130, United States

**Author notes:** Corresponding Author: Hong-Bo Pang. These authors contributed equally (Jibin Guan, Hong Guo).

## Abstract

Nucleotide-based drugs, such as antisense oligonucleotides (ASOs), have unique advantages in treating human diseases as they provide virtually unlimited ability to target any gene. However, their clinical translation faces many challenges, one of which is poor delivery to the target tissue *in vivo*. This problem is particularly evident in solid tumors. Here, we functionalized liposomes with a tumor-homing and -penetrating peptide, iRGD, as a carrier of an ASO against androgen receptor (AR) for prostate cancer treatment. The iRGD-liposomes exhibited a high loading efficiency of AR-ASO, and an efficient knockdown of AR gene products was achieved *in vitro*, including AR splice variants. *In vivo*, iRGD-liposomes significantly increased AR-ASO accumulation in the tumor tissue and decreased AR expression relative to free ASOs in prostate tumors established as subcutaneous xenografts. Similar results were obtained with intra-tibial xenografts modeling metastasis to bones, the predominant site of metastasis for prostate cancer. In treatment studies, iRGD-liposomes markedly improved the AR-ASO efficacy in suppressing the growth of both subcutaneous xenografts and intra-tibial xenografts. The inhibitory effect on tumor growth was also significantly prolonged by the delivery of the AR-ASO in the iRGD-liposomes. Meanwhile, iRGD-liposomes did not increase ASO accumulation or toxicity in healthy organs. Overall, we provide here a delivery system that can significantly increase ASO accumulation and efficacy in solid tumors. These benefits are achieved without significant side effects, providing a way to increase the antitumor efficacy of ASOs.

## Introduction

Nucleotide-based drugs, including antisense oligonucleotides (ASOs) and small interfering RNA (siRNAs), have recently gained rapid success in their translation into clinical uses [1]. Their biggest advantage lies in almost unlimited ability to target any genetically encoded material, especially those that cannot be targeted by other types of drugs, such as small molecules and antibodies [1]. This property makes them potentially useful in treating diseases with known genetic drivers, such as solid tumors [2]. In the case of prostate cancer, new therapeutic approaches are required to overcome drug resistance in the advanced stage [3] and to target bone metastasis, which occurs in 90% of lethal prostate cancer [4]. Aberrant expression and activation of androgen receptor (AR) has been recognized as the driver of progression of prostate cancer to a castration-resistant, lethal form (CRPC). Further, AR splicing variants such as variant 7 (AR-V7) have been shown to promote resistance towards small molecule AR antagonists [5,6]. While CRPC is challenging to treat with small-molecule therapeutics, AR-ASO has shown good efficacy to achieve knockdown of wild type (WT) AR and AR splicing variants, and thus yield an inhibitory effect against CRPC [7].

ASOs, as well as other nucleotide-based drugs, face several challenges *in vivo*, including nuclease degradation, renal clearance, immune responses, and nonspecific absorption by the vasculature and liver/spleen due to negative charges [8]. In solid tumors, additional transport barriers exist such as the vascular wall, multilayered stromal and tumor cells, dense extracellular matrix, and high interstitial pressure [9]. All these barriers result in poor accessibility of systemically administered ASOs to their target cells in extravascular tumor regions, which greatly limits the therapeutic efficacy of ASOs [10]. Additionally, target genes, such as AR, are often expressed in healthy tissues as well and have important roles in normal physiological functions [11,12]. Therefore, means of delivering ASOs more efficiently into solid tumors in order to avoid adverse effects and improve antitumor efficacy are needed. In metastatic cancer, the delivery would also have to include the metastatic lesions, such as bone metastases in CRPC, to be effective.

Liposomes are lipid-based nanoparticles (NPs) which have been widely used for systemic drug delivery. Multiple liposomal formulations are currently in clinical use to facilitate the delivery of small molecule drugs and siRNAs [13,14]. Liposomes encapsulate the cargo inside a lipid bilayer, which can shield the negative charges of a nucleotide and protect it from nucleases. The large size of liposomes also helps prolong the plasma half-life of the payload. To increase tumor targeting and accessibility to extravascular targets, we covalently linked a tumor-penetrating peptide, iRGD (CRGDK/RGPD/EC) onto the surface of ASO-loaded liposomes [15]. Besides selective homing to a variety of solid tumors upon systemic administration, this peptide can bring a coupled cargo, ranging from small molecules to NPs, across tumor vessels and deep into extravascular tumor tissue [15]. This vascular and tumor penetration property also applies to co-administered payloads that are not coupled to the peptide (bystander effect) [16]. The multifunctional iRGD peptide and peptides that share its RGD motif have been used in the delivery of liposomes and other nano-sized carriers to improve the drug delivery into various solid tumors, including prostate cancer and its bone metastases [17–19]. Given the central role of AR in the progression of prostate cancer, we set out here to evaluate the ability of iRGD-liposomes to enhance the delivery and efficacy of AR-ASO in mouse models of primary and metastatic prostate cancer.

## Results

### Liposome-ASO synthesis and in vitro validation

The iRGD-functionalized liposomes were synthesized as previously described with minor modifications (see Fig. 1A for schematic structure) [20,21]. Briefly, liposomes were made with two types of lipids, including polyethylene glycol (PEG) conjugated 1,2-distearoyl-sn-glycero-3-phosphoethanolamine (DSPE) (PEGylated DSPE) with or without iRGD attached to the free end of the PEG chain, and dioleoyl-3-trimethylammonium propane (DOTAP), to provide a positive charge to the interior and a charge-neutral surface to aid ASO encapsulation. An ASO against AR was acquired from IONIS therapeutics and used in the following studies. Dipalmitoyl phosphatidylcholine (DPPC) and cholesterol were added to increase the stability of liposomes and prevent premature release of liposome cargo [22,23]. Transmission electron microscopy (TEM) imaging confirmed that liposomes carrying AR-ASOs were spherical with a thin hydrated iRGD-PEG outer layer (Fig. 1B). The physicochemical properties of liposomes were measured using dynamic light scattering (DLS) (Fig. 1C). The liposomes were homogeneous in size with mean hydrodynamic diameters around 150 nm and a polydispersity index (PDI) <0.1 (Fig. 1C and Supplementary Table 1). The surface charge was negative with a zeta potential of −6.67 mV to −7.10 mV (Supplementary Table 1).

**Fig. 1.**
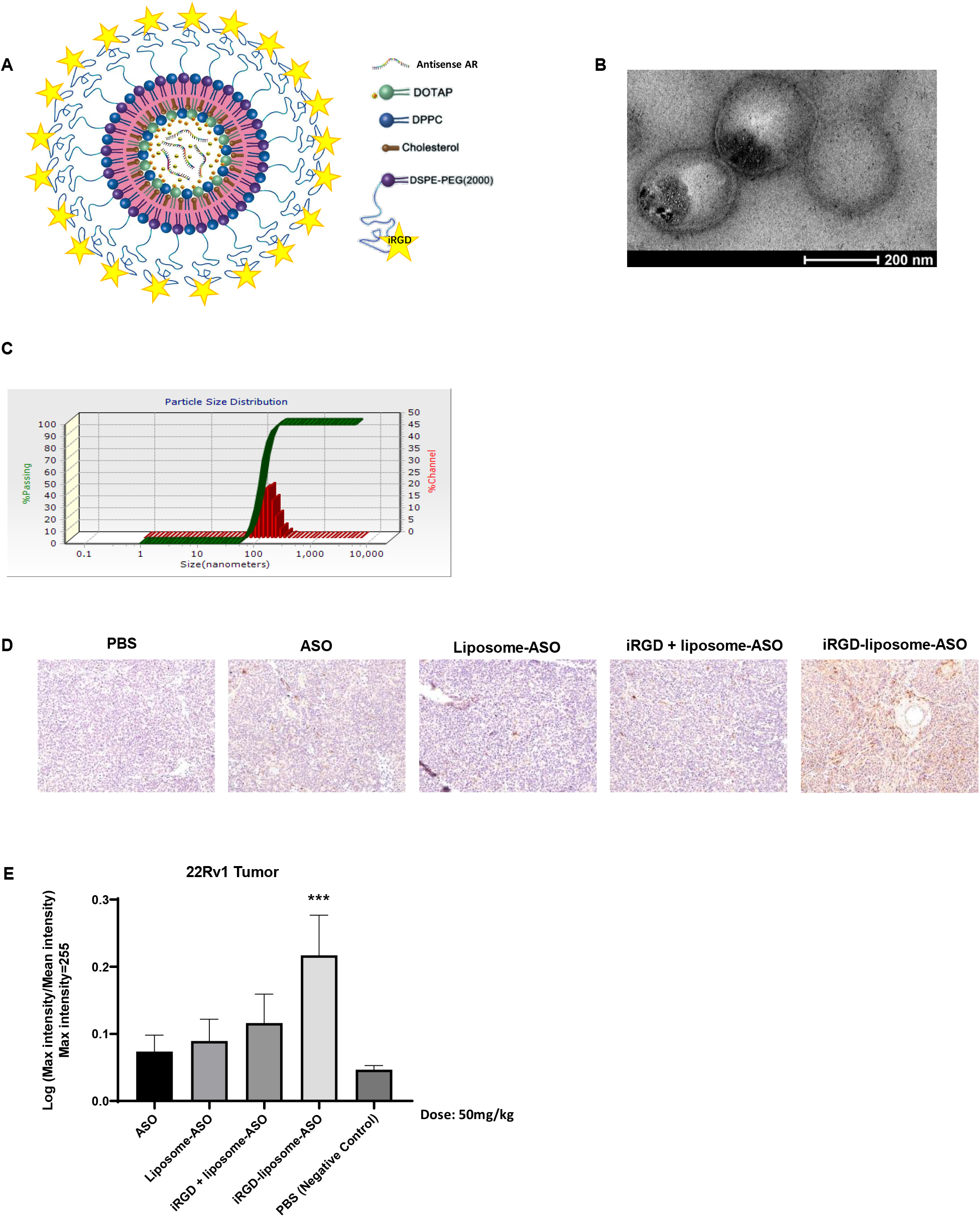
iRGD-liposome enhances homing and uptake of ASO in tumors. (A) Schematic diagram of the iRGD-liposome-ASO system showing iRGD, antisense AR, and liposomes with lipid materials (DOTAP, DPPC, Cholesterol, and DSPE-PEG2000). (B) TEM image of final constructs of iRGD-liposome-ASO showing cloudy liposomal coatings around dark cores. Scale bar: 200 nm. (C) Hydrodynamic diameter and zeta potential of iRGD-liposome-ASO. (D) Representative immunohistochemistry (IHC) of ASO staining and (E) quantification of ASO in IHC sections from tumors of 22Rv1 tumor-bearing mice. The 22Rv1 tumor-bearing mice were treated by PBS, free ASO, liposome-ASO, iRGD + liposome-ASO, and iRGD-liposome-ASO, respectively. The ASO dose was 50mg/kg. The tumors were collected after 4-day homing by different delivery systems. All experiments were performed in three mice per group. The data are represented as mean ± standard deviation (SD). ***P<0.001 vs ASO. ASO: antisense oligonucleotide; AR: androgen receptor; DOTAP: dioleoyl-3-trimethylammonium propane; DPPC: dipalmitoyl phosphatidylcholine; DSPE: 1,2-distearoyl-sn-glycero-3-phosphoethanolamine; PEG2000: polyethylene glycol 2000; PBS: phosphate-buffered saline.

An encapsulation efficiency (~8 mg/ml) of AR-ASO was obtained with the liposomes. The knockdown efficacy of this liposome-ASO formulation towards AR-FL and AR variants was studied in multiple prostate cell lines, including the 22Rv1 cell line, which displays a CRPC phenotype and harbors a structural rearrangement of the AR gene that promotes high expression of AR variants [24]. There was a significant knockdown efficiency of the liposomal formulation towards both AR-FL and AR variants in 22Rv1 cells and a reduction of AR-FL in LNCaP and VCaP cells to that of the standard transfection reagent (Lipofectamine 3000) (Fig. S1).

### iRGD-liposomes improve the AR-ASO accumulation in tumors in vivo

The major barriers for ASO delivery into solid tumors *in vivo* include low accumulation and vascular/tissue penetration in tumors. To overcome these limitations, we used ASO liposomes that incorporate an iRGD surface coating. We first optimized various aspects of particles and their use (size, dose, and dosing scheme) using 4T1 mouse breast tumor model. Two different sizes of iRGD-liposomes were tested and their ASO encapsulation efficiency was listed in Supplementary Table S1. After intravenous injection, iRGD-liposomes with a smaller size (diameter ~150 nm) achieved a higher amount of ASO accumulation in the tumor than larger iRGD-liposomes (Fig. S2 A-B). Based on these findings, we thus chose the 150-nm iRGD-liposome formulation for further studies. Varying the dose of ASO loaded in the iRGD-liposomes showed that ASO accumulation in the tumor increased in a dose dependent manner (Fig. S2 C-D). Comparison of iRGD-liposomes, non-targeted liposomes, and co-administration of non-targeted liposomes with free iRGD peptide [25], showed that iRGD-liposomes resulted in greater ASO accumulation in the tumor than the other two formulations (Fig. S2 C-D). Finally, we investigated ASO accumulation after a longer time of circulation (4 days post injection vs 4 hours). The iRGD-liposome group still exhibited the highest spreading and retention of ASOs in the tumor (Fig. S2 E-F). The ASO signals were dispersed across the tumor, rather than confined in the perivascular regions (Fig. S2G). An AR-positive CRPC tumor model, 22Rv1 [6] gave similar results (Fig. 1 D-E).

The biodistribution in healthy organs of ASOs delivered by the different formulations was also investigated at different time points after systemic administration. A substantial accumulation of ASOs was observed in the liver and kidney, while no or much less in other organs. No differences in healthy organ accumulation were observed across different formulations, liposomal sizes, ASO dosages and time points (Fig. S3 A-C). This result suggests that iRGD-liposomes mainly improve the ASO accumulation and vascular penetration in the tumor, with minimal effects on its accumulation in healthy organs.

### Pharmacokinetics and biocompatibility of iRGD-liposome-ASO in vivo

Pharmacokinetic parameters for the liposomal ASO formulations were calculated using a noncompartmental model based on the plasma AR-ASO concentrations (Supplementary Table S2 and Fig. S4A). Compared to free ASOs, liposomal formulations improved ASO-bioavailability (F%), the area under the plasma concentration curve (AUC), and plasma half-life (T_1/2_), while iRGD yielded little additional effect. The levels of AR-ASO in plasma were low in all groups 24 h after injection. This result suggests that iRGD mainly improves the tumor accumulation of liposomal ASOs while having little effect on their pharmacokinetic profile.

To evaluate whether liposomal formulations increase the toxicity of ASOs, we assessed the *in vivo* biological safety by measuring blood toxicity indicators (liver enzymes aspartate transaminase (AST) and alanine transaminase (ALT)) in tumor-bearing mice after injecting the highest dosage of ASOs used in this study. There was little difference between liposomal groups and free ASOs in terms of AST/ALT levels (Fig. S4B).

### iRGD-liposomes increase the therapeutic efficacy of AR-ASO against subcutaneous prostate cancer xenografts

Using the CRPC induced by transplanting 22Rv1 cells, we tested different ASO dosages and injection frequency. Compared to the PBS group, ASO treatment in all groups showed antitumor efficacy of various degrees (Fig. S5 A-C). iRGD-liposomes, when carrying an equivalent amount of ASOs, exhibited a much stronger inhibitory effect towards tumor growth than the free ASO group regardless of injection frequency. Quantification of ASO (Fig. S5 D) and AR protein expression (Fig. S5 E) in the tumor by immunohistochemistry (IHC) showed a significant increase of ASO and reduction in AR protein, which aligns with the tumor treatment results. Lowering the ASO amount in the iRGD-liposomes, however, greatly reduced their efficacy (Fig. S5 F and G). The body weight remained consistent among all groups during the two-week treatment, further supporting that iRGD-liposomes result in little toxicity compared to free ASOs (Fig. S5H).

To evaluate the contribution of iRGD to liposome efficacy, we compared liposome-ASO and iRGD-liposome-ASO using the 22Rv1 model. The ASO dose was 50mg/kg and the injections were given every four days. Liposomal encapsulation significantly enhanced the ASO ability to suppress tumor growth, and iRGD gave further improvement (Fig. 2 A-B, Fig. S6A). IHC staining showed increased ASO staining in the tumor and reduced AR expression that paralleled with the increase in antitumor efficacy of the liposomal formulations (Fig. 2 C-F). TUNEL (terminal deoxynucleotidyl transferase dUTP nick end labeling) assay to assess the apoptosis levels in tumor and healthy tissues showed that iRGD-liposome-ASO induced the highest level of apoptosis in the tumor, followed by liposome-ASO and free ASO, while their effect on TUNEL staining in healthy tissues did not differ from the PBS control (Fig. 2G, Fig. S6B). Collectively, these results demonstrate that iRGD-liposomes increase the ASO accumulation and suppress AR expression selectively in the tumor, resulting in a higher cellular apoptosis and a stronger tumor inhibition. Another prostate tumor model, VCaP, which was initially isolated from human bone metastasis with a high FL-AR expression due to AR gene amplification [26], gave similar results: iRGD-liposome-ASO exhibited the strongest tumor inhibitory effect based on tumor volume and weight (Fig. 2 H-I).

**Fig. 2.**
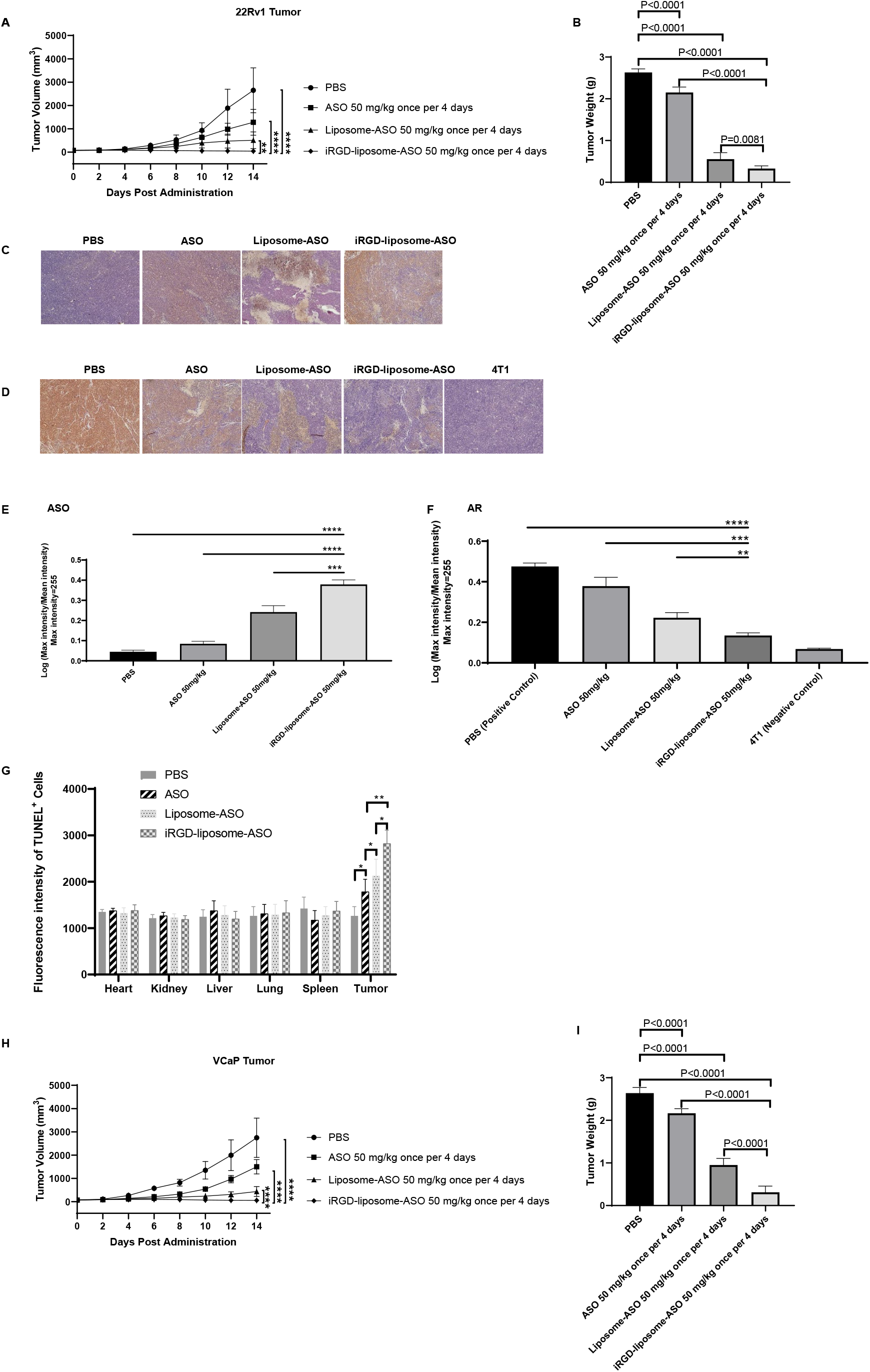
Short-term therapeutic efficacy of tumor cell-targeting liposomes loaded with ASO (iRGD-liposome-ASO) in CRPC tumors. (A) Tumor volume growth curves and (B) average tumor weight of 22Rv1 tumor-bearing mice. The 22Rv1 tumor-bearing mice were treated with PBS, free ASO, liposome-ASO, and iRGD-liposome-ASO, respectively. The ASO dose was 50mg/kg and the injections were given every four days for two weeks. (C) Representative images of ASO staining and (D) AR staining by IHC in tumors of 22Rv1 tumor-bearing mice. The ASO dose was 50mg/kg and the injections were given every four days for two weeks. AR staining in 4T1 tumor serves as the negative control. (E) Quantification of ASO and (F) AR density in IHC sections from tumors of 22Rv1 tumor-bearing mice. (G) Quantification of immunofluorescent signals of TUNEL positive cells showing apoptotic cells in tumor and other tissues of 22Rv1-bearing mice with different treatments. The ASO dose was 50mg/kg and the injections were given every four days for two weeks. (H) Tumor volume growth curves of VCaP tumor-bearing mice treated with different treatments for two weeks. (I) Average tumor weight of mice bearing VCaP subcutaneous xenografts at the end of treatment. The VCaP tumor-bearing mice were treated with PBS, free ASO, liposome-ASO, and iRGD-liposome-ASO, respectively. The ASO dose was 50mg/kg and the injections were given every four days for two weeks. The density of ASO and AR expression as well as fluorescence density of TUNEL assay were analyzed by ImageJ. All experiments were performed in 5-6 mice per group. The data are presented as the mean ± standard deviation (SD). Statistical significance was calculated using Student’s t test. The data are represented as mean ± standard deviation (SD). *P<0.05, ***P<0.001, **** P<0.0001 vs PBS controls.

In addition to the treatment described above, where the mice were sacrificed after two weeks of treatment, we also performed a study in which the mice were observed for 4 months after the end of the treatment. The tumor sizes of iRGD-liposome-ASO group remained negligible during the 4-month observation period, whereas the tumors in the PBS and free ASO groups reached the endpoint sizes within three weeks (Fig. 3 A-C).

**Fig. 3.**
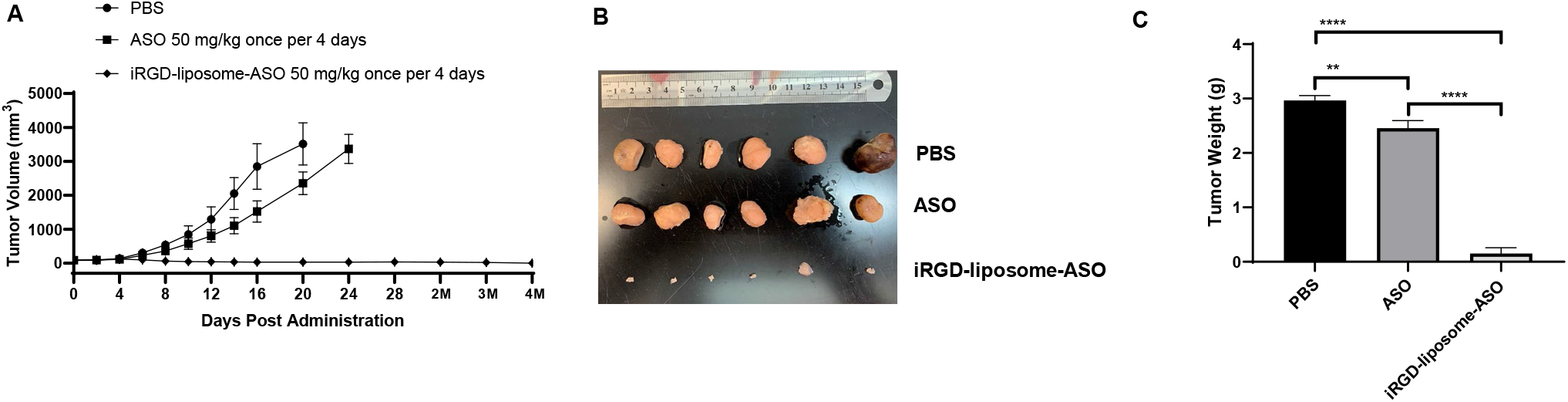
Long-term therapeutic efficacy of iRGD-liposome-ASO in CRPC tumors. The 22Rv1 tumor-bearing mice were treated with PBS, free ASO, and iRGD-liposome-ASO, respectively. The ASO dose was 50mg/kg and the injections were given every four days for two weeks. The long-term effect of iRGD-liposome-ASO are presented after 4 months of treatment by the growth curve of tumor volume (A), images of tumor (B), and the tumor weight by the end of observation (C). All experiments were performed in 5-6 mice per group. The data are represented as mean ± standard deviation (SD). *P<0.05, ** P<0.01, ***P<0.001, **** P<0.0001.

### iRGD-liposome-ASO inhibits the growth of prostate cancer bone metastasis

Bone is the primary metastasis site of prostate cancer. We used a CRPC bone metastasis model by intratibial injection of enzalutamide-resistant CWR-R1 cells [27] to study the effect of liposomal ASO delivery on metastases. Liposomal ASO formulations significantly increased the ASO accumulation in cancerous bone compared to free ASO (Fig. S7 A-B). There was little difference in ASO amounts across these groups in healthy bones and control organs (Fig. S7 A, C). In a two-week treatment study, the iRGD-liposome-ASO exhibited the highest tumor-inhibitory effect based on tumor weight (Fig. 4 A-B). Based in IHC staining at the end of the treatment, iRGD-liposomes had increased the ASO levels in the cancerous bone, but not healthy bone, with corresponding suppression of AR expression (Fig. 4 C-F).

**Fig. 4.**
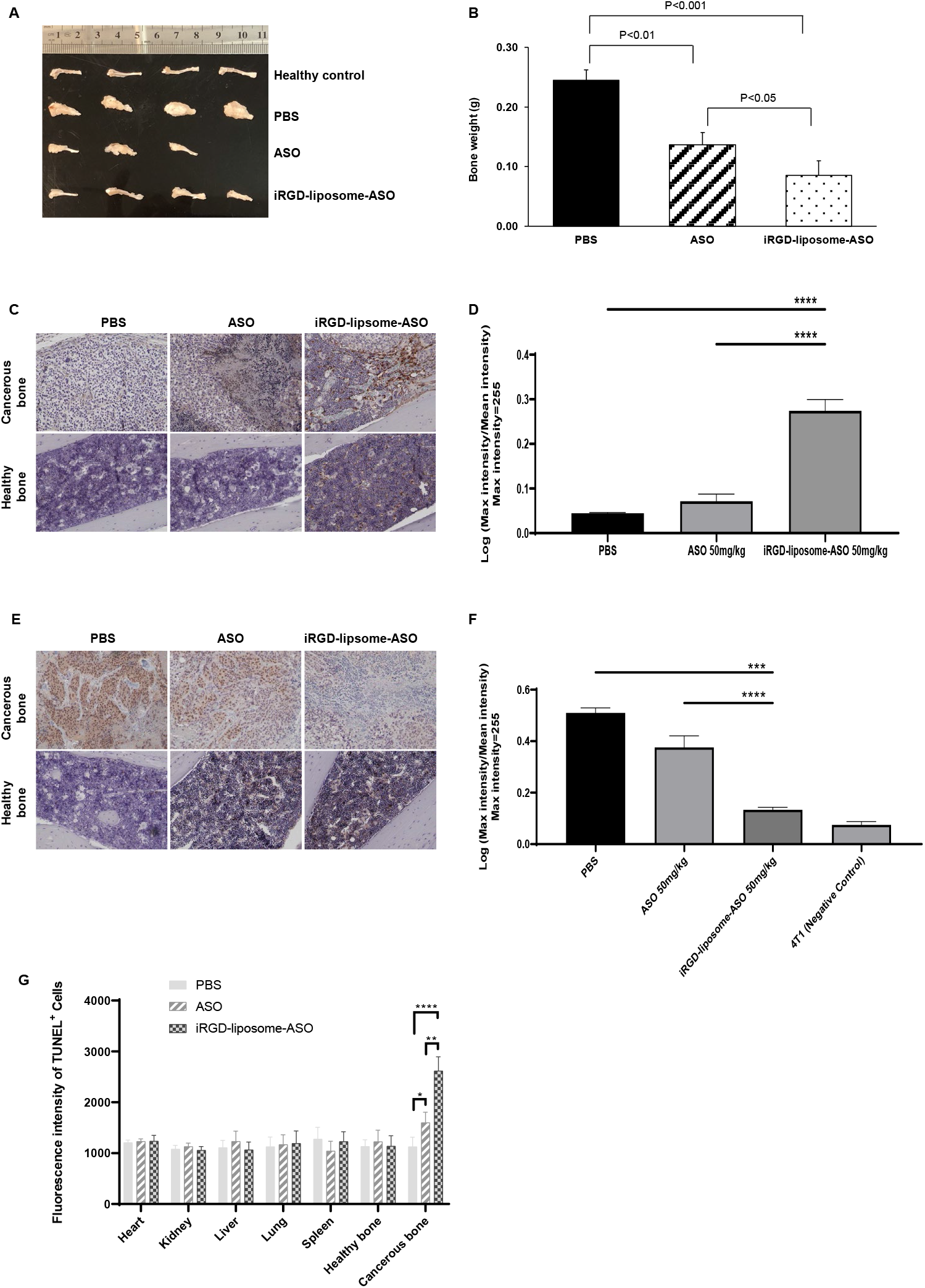
The tumor cell-targeting liposomes enhance homing and therapeutic efficacy in the CRPC bone metastasis model. The mice bearing bone metastasis were treated with PBS, free ASO, and iRGD-liposome-ASO, respectively. The ASO dose was 50mg/kg and the injections were given every 2 days for two weeks. (A) Images of bones and (B) average bone weight of mice from each group after 2 weeks of treatments by PBS, ASO, and iRGD-liposome-ASO, with a 50 mg/kg dosage of ASO every 2 days, respectively. (C) Representative images of ASO staining by IHC and (D) quantification of ASO density in healthy and metastatic bones after 2-wk treatments. (E) Representative IHC images of AR staining and (F) quantification of AR density in bones after 2-wk treatments. (G) Quantification of immunofluorescent signals of TUNEL positive cells showing apoptotic cells in tumor and other tissues of mice bearing bone metastasis with a 50mg/kg dosage of ASO every 2 days for two weeks. All experiments were performed in 3-4 mice per group. The density of ASO and AR expression were analyzed by ImageJ. The data are represented as mean ± standard deviation (SD). * P<0.05, ** P<0.01, *** P<0.001, **** P<0.0001.

There was no difference in ASO accumulation in healthy organs after two-week treatment between the iRGD-liposome-ASO and free ASO groups (Fig. S7D). No significant changes in body weight were seen during the treatment period (Fig. S7E). We also assessed the apoptosis levels in metastatic bones and healthy bone/tissues. Similar to primary tumors, iRGD-liposomes enhanced the ASO ability to induce apoptosis selectively in metastatic bone but not elsewhere (Fig. 4G, Fig. S7F). Together, these results demonstrate that iRGD-liposomes enhance the antitumor efficacy of AR-ASOs against both primary prostate tumors and bone metastases by increasing the tumor delivery of ASO.

## Discussion

In this study, we used iRGD-functionalized liposomes to improve the tumor delivery and antitumor efficacy of an ASO against AR gene and its splicing variants for CRPC treatment. The liposomes showed a satisfactory encapsulation efficiency of AR-ASO, and liposomal delivery achieved an effective knockdown of AR gene expression *in vitro* and *in vivo*. In multiple prostate tumor models, iRGD and liposomes showed a synergistic effect to increase ASO accumulation in the tumor, but not elsewhere, compared to free AR-ASO. IHC analysis also showed that the ASO was distributed broadly in the tumor tissue, rather than confined to areas near the blood vessels. In addition to reduced AR expression, the increased ASO delivery led to increased apoptosis level, and thus more effective suppression of tumor growth in multiple CRPC models. Besides subcutaneous tumor xenografts, we also validated the benefit of iRGD-liposomal delivery in a bone metastatic prostate cancer. Together, we showcase here a tumor-targeted delivery system for an ASO that may be applicable to the delivery of nucleotide-based drugs in general, allowing improvement in their efficacy.

One of the major challenges in the *in vivo* applications of ASOs and other nucleotide drugs, especially for solid tumors, is the delivery. Lipid-based NPs have been the mainstream carriers to solve this problem to some extent, by protecting the nucleotides and shielding their negative charges. However, compared to siRNAs and even CRISPR-Cas9, much fewer studies have focused on ASO drugs [28,29]. Here, our results demonstrated the feasibility of liposomal system in improving ASO delivery. Pharmacokinetic analysis showed that liposomes prolong the plasma half-life and AUC of ASOs, and toxicity/IHC studies indicated that liposomal formulations do not induce additional toxicity.

One feature of nucleotide-based drugs is that they are usually well tolerated, at least compared to common chemotherapy drugs, due to the high target specificity of ASOs [7]. Thus, the main challenge lies in the inefficient entry of ASOs into tumor tissue, rather than nonspecific accumulation and the resulting adverse effects in healthy organs. Liposomes and other NPs have been long thought to enhance drug delivery into solid tumors via passive diffusion through intercellular gaps (enhanced permeation and retention (EPR) effect) [30]. However, recent studies have revealed that this route only accounts for a small fraction of nanoparticle entry into solid tumors, and their penetration distance from the vasculature is limited due to additional transport barriers in the stroma [25,31]. Therefore, we employed iRGD, which triggers an active trans-vascular, trans-tissue transport process that delivers cargo deep into tumor tissue [15,32]. Our results show that while liposomal encapsulation alone enhances ASO accumulation in tumors, iRGD functionalization of the liposomes increases it further. Optimizing formulation of iRGD-liposomes increased the accumulation of ASO in the tumor, while not affecting ASO biodistribution in healthy tissues, further broadening the therapeutic window.

Nucleotide-based drugs such as ASOs have unique advantages in therapeutic targeting of genetic alterations, such as those in AR. The progression of prostate cancer to a CRPC phenotype is driven by aberrant expression and/or activation of AR gene and its related signaling events. These changes occur in response to treatment with AR-targeted compounds such as enzalutamide and abiraterone [33–36], which prolong the survival of CRPC patients [6,37,38]. The changes in the *AR* gene include amplification, mutation, structural rearrangement in 70-90% of CRPC tumors [39,40]. Further variability is caused by changes in the expression of AR variants resulting from altered mRNA splicing and polyadenylation [41]. The ASO used in this study has been previously shown to effectively knock down both the wild-type AR gene and splicing variants, leading to the suppression of CRPC tumors resistant to AR antagonists [6]. However, a high dosage and injection frequency were required to obtain a clinical benefit [28]. Moreover, AR is expressed in many healthy tissues (e.g. bone, muscle, and brain), and has important functions in normal physiology [42–44]. Therefore, it is desirable to increase ASO delivery into the tumor, while minimizing ASO accumulation in healthy organs. Our results show that an iRGD-liposome system can achieve these two goals. Moreover, at an equivalent amount of ASOs, iRGD-liposome-ASO gave the same or stronger antitumor effect compared to free AR-ASO with fewer injections. Intriguingly, after the last administration, iRGD-liposome-ASO resulted in long-term suppression of CRPC growth in the 22Rv1 model. Thus, iRGD-liposomes seem a promising way of accelerating the clinical translation of ASO drugs as alternatives for CRPC treatment.

As is common knowledge, it is metastasis, rather than primary tumors, that makes cancer lethal [4]. Thus, we also investigated whether AR-ASO has therapeutic effect against bone metastases (the primary metastatic site of prostate cancer), and whether iRGD-liposomes would enhance the ASO delivery to metastatic bones. NP-based carriers, including liposomes, have been used for drug delivery to bone metastases using passive and active targeting [19]. Passively targeted NPs with a size smaller than bone marrow fenestrations can pass through the capillaries and accumulate in bone marrow [45]. Active ligand-directed targeting has also been explored, some all the way to clinical trials [46,47]. A notable example is bisphosphonate [48], which specifically binds to bone hydroxyapatite matrix [49], and has been applied in cancer-associated bone diseases [50–52]. Unfortunately, bisphosphonate-conjugated drug accumulates in all bones and primarily targets osteoclasts rather than tumor cells [53,54]. RGD peptides have also been used to target bone metastases [55], and iRGD was originally identified by screening for peptides that accumulate in bone metastases of prostate cancer [15], which prompted us to investigate whether iRGD in ASO targeting to metastases. Our results show that iRGD-liposomes also enhance the AR-ASO accumulation in metastatic bones but not to healthy bones. Encapsulation AR-ASO into iRGD-liposomes strongly inhibited the growth of bone metastases, significantly exceeding the effect of the ASO alone.

Taken together, the results here presented show that iRGD-liposomes provide an effective delivery system for AR-ASOs, both in primary tumor and bone metastases, and that this translates to an improved antitumor efficacy. As previously reported for iRGD functionalized liposomes with other payloads, this strategy may be applicable to other types of nucleotide-based therapeutics, and a variety of other solid tumor types and metastasis sites.

## Materials & methods

### Chemicals and reagents

Chemically modified AR-ASO (sense strand, 5′TGATTTAATGGTTGCA3′) and Cy3-conjugated ASO (5′GCATTCTAATAGCAGC3′) were obtained from Ionis Pharmaceuticals (Carlsbad, CA). FAM-cysteine-iRGD was purchased from LifeTein (Somerset, NJ). DSPE-PEG2000, DPPC, DOTAP, and cholesterol were obtained from Avanti Polar Lipid (Alabaster, Alabama). DSPE-PEG2000-MAL was from Nanosoft Polymers (Winston-Salem, NC). The Lipofectamine 3000 transfection kit was from Invitrogen (Carlsbad, CA). Dulbecco’s modified Eagle’s medium (DMEM), RPMI-1640 Medium, and trypsin were from Sigma-Aldrich (St Louis, MO). All reagents and compounds were used without further purification or modification.

### Preparation of iRGD-liposome-ASO

Lipids were weighed and dissolved with chloroform. The total lipids were composed under a proper molar ratio (DPPC, DSPE-PEG-Mal/OCH_3_, DPTAP, cholesterol as 15: 2: 3: 10). The chloroform was removed by rotary evaporation and under nitrogen blowing. The film was hydrated with 10 mL ASO HBS (10 mM pH 7.4) solution containing 10% sucrose by using the rotavap under 40 °C water bath condition but at normal atmosphere for around two hours. The primary liposomes were extruded by using membrane with a pore size of 1 μm, then frozen to −80 °C and further lyophilized. The liposomes were hydrated with HBS following the dry-freezing procedure, and before the extrusion. To compare the homing efficiency of two particle sizes (100 nm and 200 nm), the hydrated liposomes were extruded by using membranes with pore size of 0.1 μm and 0.2 μm, respectively. Further, proper molar of FAM-cysteine-iRGD was added into the solution to conjugate to the maleimide on the surface of the liposomes. In the end, the ASO-liposome-iRGD was dialyzed with a 50kDa kit to obtain the final product. For the evaluation of ASO content in the liposomes, the nucleic acid concentration after damaging the lipid bilayers with 10% Triton was determined by Nanodrop.

### Characterization of iRGD-liposome-ASO systems

The average size and size distributions of liposomes were determined by dynamic light scattering (DLS). DLS measurements were performed using a NanoFlex Particle Analyzer (Microtrac, Montgomeryville, PA). Autocorrelation functions were analyzed by the cumulants method (fitting a single exponential to the correlation function to obtain the mean size and the PDI and the CONTIN routine (fitting a multiple exponential to the correlation function to obtain the distribution of particle sizes). All measurements were performed at a 90° angle. The Zeta potential of the liposomes were measured by Malvern ZETASIZER Nano ZS. The morphology images of the liposomes were acquired using a TECNAI G2 Spirit BIOTWIN transmission electron microscope.

### Cell culture

Human 22Rv1, LNCaP, and VCaP cells were purchased from American Type Culture Collection (ATCC, Manassas, VA). 22Rv1 and LNCaP cells were cultured in RPMI-1640 medium containing 50 U per mL streptomycin, 100 U per mL penicillin, and 10% fetal bovine serum (FBS, Thermo Fisher Scientific, Waltham, MA). VCaP cells were cultured with DMEM medium containing 50 U per mL streptomycin, 100 U per mL penicillin, and 10% FBS. The cells were cultured at 37°C in a humidified incubator with 5% CO_2_. All the cell cultures were maintained in 25 cm^2^, 75 cm^2^ cell culture flasks, or 10 cm culture dishes for use.

### Western blotting

The protein extracts from the cells (40 μg) were subjected to electrophoresis in SDS-PAGE on 4–15% precast protein gels (Bio-Rad, Hercules, CA), separated and transferred to polyvinylidene fluoride (PVDF, 0.2 μm pore size) membranes. The membranes were probed with primary antibodies to the androgen receptor (GeneTex, Irvine, CA) and β-actin (Thermo Fisher Scientific) overnight at 4 °C. After washing, the membranes were incubated with the secondary antibodies IRDye® 680RD donkey anti-rabbit IgG and IRDye® 800CW donkey anti-mouse IgG (Li-COR, Lincoln, NE). The bands were visualized by scanning the fluorescence signal via the Odyssey® CLx imaging system (LI-COR). Then the photographic images with blots were scanned and quantified by the software of the system for the comparison of the fluorescence value of the blots.

### Immunohistochemistry (IHC) staining

Immunohistochemical staining was performed on the paraffin-embedded sections. In brief, after deparaffinization and rehydration, all sections were incubated with 0.3% H_2_O_2_ solution for blockade of endogenous peroxidase activity. The sections were then blocked with 5% donkey serum blocking solution (with 0.1% triton X100), followed by incubation with primary antibodies overnight at 4°C. The primary antibodies included monoclonal rabbit anti-ASO (Ionis, Carlsbad, CA) and polyclonal rabbit anti-AR (PA1-110, Invitrogen). The bound antibodies were detected using ImmPACT™ DAB Peroxidase (HRP) Substrate Kit (Vector Labs, Burlingame, CA) followed by hematoxylin counterstaining. The CD31 antibody (MA1-40074, Thermo Fisher Scientific) was used for the co-staining of blood vessels with ASO. The bounded antibodies were detected using Vector AEC peroxidase substrate kit (Vector Labs). Images of randomly selected areas from each section were collected by the microscope. Three views per tissue were captured from each section for semi-quantification by ImageJ software according to the HRP-DAB signal.

### Immunofluorescence (IF) staining

Immunofluorescence staining was performed on the paraffin-embedded sections. In brief, after deparaffinization and rehydration, all sections were treated with PBS containing 1% BSA and 0.1% Triton X100 (blocking buffer) at room temperature (RT) for 1 h. The sections were washed three times with PBS and then incubated with primary antibodies anti-ASO (Ionis Pharmaceuticals. Carlsbad, CA) and anti-CD31 (BD Bioscience, San Jose, CA) with a 1:200 dilution in blocking buffer at 4 °C overnight, followed by the appropriate secondary antibodies diluted (1:200) in blocking buffer at RT for 1 h. After washing with PBS, sections were mounted in DAPI-containing mounting medium (Vector Laboratories, Burlingame, CA) with a coverslip and examined under fluorescence microscope EVOS M5000 (Thermo Fisher Scientific).

### Animals

All animal studies were carried out in compliance with the National Institutes of Health guidelines and an approved protocol from University of Minnesota Animal Care and Use Committee. The animals were housed in a specific pathogen-free facility with free access to food and water at the Research Animal Resources (RAR) facility of the University of Minnesota.

Female BALB/c mice (6~7 weeks old; weight at 20–25 g) and Male athymic nude mice (6~7 weeks old; weight at 20–25 g) were purchased from Charles River and Envigo (Indianapolis, IN), respectively. The 4T1 orthotopic tumor model was generated by the injection of 4 million of 4T1 cells into the breast of each female mouse. The tumor would be ready after two weeks of injection. The xenograft 22Rv1 and VCaP tumor models were established by subcutaneously injecting 5 × 10^6^ 22Rv1 / 8 × 10^6^ VCaP cells in normal saline into the lower flank of the male mice, respectively. The tumor diameters were measured with calipers every two days, and tumor volumes were calculated as V_tumor_ = width^2^ × length/2. After 2-3 weeks (22Rv1) or 5-6 weeks post-tumor inoculation, tumor size reached around 5 mm in diameter.

Bone Metastasis Model of CRPC was generated via intratibial injection of luciferase-expressing prostate cancer cells R1-EnzR (enzalutamide-resistant CWR-R1 cells), which were kindly provided by Dr. Aaron LeBeau at University of Minnesota. In brief, 4~6 weeks old male BALB/c nude mice (Envigo) were anesthetized by isoflurane with the proximal end of the left tibia exposed with the knee maintained in a flexed position. In order to create a hole/passage, a 26-gauge needle was inserted into the joint surface of the tibia through the patellar tendon and then down into the bone marrow cavity with a drilling motion. Subsequently, 10 ul of cell solution (1 × 10^6^ cells in PBS) was slowly injected into the tibia bone marrow cavity. Animals were monitored daily for body weight and signs of tumor burden.

### *In vivo* pharmacokinetics

Wild-type C57BL/6 male mice (6~7 weeks old; weight at 20–25 g) were deprived of food overnight but having free access to water. The mice were randomly divided into five groups and were injected via lateral tail vein with the formulation of ASO solution, liposome-ASO and with equivalent AR-ASO (with 1% CY3-conjugated ASO) dose (50 mg/kg), respectively. Approximately 100 μL of blood was collected into tubes via venous plexus at 5, 10, 20, 30, 45, 90 min and, 2, 6, 12, 24, 48 and 72 h, then centrifuged at 13200 rpm for 10 min to obtain plasma. Then the plasma samples were frozen at −80 °C for fluorescent analysis. CY3-ASO standards were prepared by blending working standard solutions to gain effective concentration levels of 0.5, 1, 2.5, 5, 10, 25, 50, 100, 250, 500 ng/mL with plasma (50 μL). Fluorescence was analyzed at 540/570 nm wavelength using a microplate reader (BioTek, Winooski, VT).

### *In vivo* targeting & antitumor assay

The homing procedure was performed in 4T1 tumor-bearing mice by administering PBS, ASO, ASO-liposome, and ASO-liposome-iRGD *via* a one-time tail vein intravenous injection at 10/50 mg/kg dosage. After 2 hours /4 hours /4 days, mice were euthanized and the tissues were harvested.

The treatments of 22Rv1 and VCaP xenograft tumor-bearing mice were initiated at a 48 hours interval for two weeks and the tumor growth was monitored for 4 months (22Rv1 model). The mice were randomly divided into groups (*n* = 6 per group) receiving different injections as follows: (1) PBS, (2) free ASO in PBS, (3) ASO-liposome, (4) ASO-liposome-iRGD. The dosage for the mice were 20 and 50 mg/kg by two different injection frequencies (every two days or four days). Tumor volume and body weight were recorded. The obtained tumors were accurately weighed. All the harvested tissues were fixed, processed, and embedded in paraffin for use.

### TUNEL staining

The TUNEL staining was performed using an In Situ Cell Death Detection Kit (Roche Diagnostics GmbH, Mannheim, Germany). The paraffin sections were stained with TUNEL reaction mixture for 1 h at 37°C and then mounted using DAPI containing mount medium. The images were captured by using fluorescence microscopy EVOS M5000.

### Serum Biochemical Toxicity Analysis

Serum aminotransferase quantification was performed to determine toxicity effect of different formulations of AR-ASO on the liver. Blood samples were collected by cardiac puncture after treatment. Serum aspartate aminotransferase (AST) and alanine aminotransferase (ALT) levels were measured using assay kits from Stanbio (Boerne, TX).

### Statistical analysis

All statistical analyses in this paper were presented as mean ± SD. Statistical analysis was performed using a one-way analysis of variance (ANOVA) or Student’s t-test, where appropriate to assess differences among groups. The differences were considered significant at *P < 0.05, **P < 0.01, ***P < 0.001 and ****P < 0.0001.

## Author Contributions

H-B.P., E.R., and P.S. designed the project. Y.L. and S.M.D. helped design and performed bone metastasis studies. J.G., H.G., Y. W., T.T., and Y.W. carried out the rest of the study. J.G., H.G. and H-B.P. wrote the manuscript.

## Acknowledgements

We thank Dr. Michael J. Sailor, Dr. Aaron LeBeau, Dr. William Elmquist, Dr. Stephen Hecht and Dr. Yupeng Li for their kind help on reagents and instrument. Research reported in this publication was supported by grants from the National Institute of Health (R01CA214550, R01GM133885, R21EB022652) and the State of Minnesota (MNP#19.08). The content is solely the responsibility of the authors and does not necessarily represent the official views of the National Institutes of Health. Portions of this work were conducted in the Minnesota Nano Center, which is supported by the National Science Foundation through the National Nanotechnology Coordinated Infrastructure (NNCI) under Award Number ECCS-2025124.

## Graphical Abstract

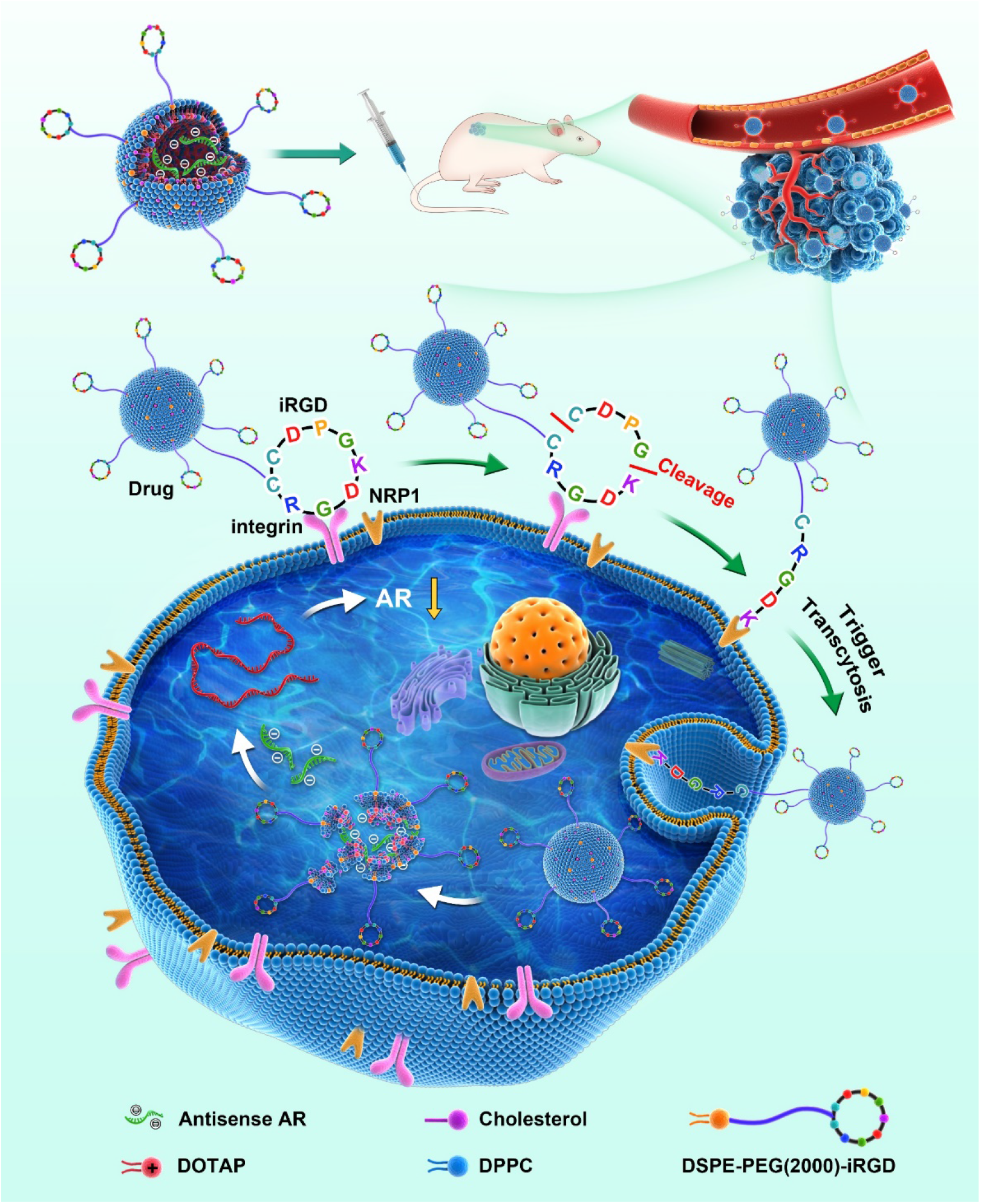

## References

[1] C.X. Li, A. Parker, E. Menocal, S. Xiang, L. Borodyansky, J.H. Fruehauf, Delivery of RNA interference, Cell Cycle. 5 (2006) 2103–2109. https://doi.org/10.4161/cc.5.18.3192.

[2] J.E. Zuckerman, I. Gritli, A. Tolcher, J.D. Heidel, D. Lim, R. Morgan, Correlating animal and human phase Ia / Ib clinical data with CALAA-01, a targeted, polymer-based nanoparticle containing siRNA, 111 (2014). https://doi.org/10.1073/pnas.1411393111.

[3] T.M.S. Amaral, D. Macedo, I. Fernandes, L. Costa, Castration-Resistant Prostate Cancer: Mechanisms, Targets, and Treatment, Prostate Cancer. 2012 (2012) 1–11. https://doi.org/10.1155/2012/327253.

[4] L. Bubendorf, A. Schöpfer, U. Wagner, G. Sauter, H. Moch, N. Willi, T.C. Gasser, M.J. Mihatsch, Metastatic patterns of prostate cancer: An autopsy study of 1,589 patients, Hum. Pathol. 31 (2000) 578–583. https://doi.org/10.1053/hp.2000.6698.

[5] H.I. Scher, C.L. Sawyers, Biology of progressive, castration-resistant prostate cancer: Directed therapies targeting the androgen-receptor signaling axis, J. Clin. Oncol. 23 (2005) 8253–8261. https://doi.org/10.1200/JCO.2005.03.4777.

[6] Y. Li, S.C. Chan, L.J. Brand, T.H. Hwang, K.A.T. Silverstein, S.M. Dehm, Androgen receptor splice variants mediate enzalutamide resistance in castration-resistant prostate cancer cell lines, Cancer Res. 73 (2013) 483–489. https://doi.org/10.1158/0008-5472.CAN-12-3630.

[7] Y. Yamamoto, Y. Loriot, E. Beraldi, F. Zhang, A.W. Wyatt, N. Al Nakouzi, F. Mo, T. Zhou, Y. Kim, B.P. Monia, A.R. MacLeod, L. Fazli, Y. Wang, C.C. Collins, A. Zoubeidi, M. Gleave, Generation 2.5 antisense oligonucleotides targeting the androgen receptor and its splice variants suppress enzalutamide-resistant prostate cancer cell growth, Clin. Cancer Res. 21 (2015) 1675–1687. https://doi.org/10.1158/1078-0432.CCR-14-1108.

[8] P.P. Seth, M. Tanowitz, C. Frank Bennett, Selective tissue targeting of synthetic nucleic acid drugs, J. Clin. Invest. 129 (2019) 915–925. https://doi.org/10.1172/JCI125228.

[9] C. Heldin, K. Rubin, K. Pietras, A. Östman, HIGH INTERSTITIAL FLUID PRESSURE — AN OBSTACLE IN CANCER THERAPY, 4 (2004) 806–813. https://doi.org/10.1038/nrc1456.

[10] M.W. Dewhirst, T.W. Secomb, Transport of drugs from blood vessels to tumour tissue, Nat. Publ. Gr. 17 (2017) 738–750. https://doi.org/10.1038/nrc.2017.93.

[11] D.K. Lee, C. Chang, Endocrine mechanisms of disease: Expression and degradation of androgen receptor: Mechanism and clinical implication, J. Clin. Endocrinol. Metab. 88 (2003) 4043–4054. https://doi.org/10.1210/jc.2003-030261.

[12] K. De Gendt, G. Verhoeven, Tissue- and cell-specific functions of the androgen receptor revealed through conditional knockout models in mice, Mol. Cell. Endocrinol. 352 (2012) 13–25. https://doi.org/10.1016/j.mce.2011.08.008.

[13] Y.C. Barenholz, Doxil ® — The first FDA-approved nano-drug : Lessons learned, J. Control. Release. 160 (2012) 117–134. https://doi.org/10.1016/j.jconrel.2012.03.020.

[14] D. Adams, A. Gonzalez-Duarte, W.D. O’Riordan, C.-C. Yang, M. Ueda, A. V. Kristen, I. Tournev, H.H. Schmidt, T. Coelho, J.L. Berk, K.-P. Lin, G. Vita, S. Attarian, V. Planté-Bordeneuve, M.M. Mezei, J.M. Campistol, J. Buades, T.H. Brannagan, B.J. Kim, J. Oh, Y. Parman, Y. Sekijima, P.N. Hawkins, S.D. Solomon, M. Polydefkis, P.J. Dyck, P.J. Gandhi, S. Goyal, J. Chen, A.L. Strahs, S. V. Nochur, M.T. Sweetser, P.P. Garg, A.K. Vaishnaw, J.A. Gollob, O.B. Suhr, Patisiran, an RNAi Therapeutic, for Hereditary Transthyretin Amyloidosis, N. Engl. J. Med. 379 (2018) 11–21. https://doi.org/10.1056/nejmoa1716153.

[15] K.N. Sugahara, T. Teesalu, P.P. Karmali, V.R. Kotamraju, L. Agemy, O.M. Girard, D. Hanahan, R.F. Mattrey, E. Ruoslahti, Tissue-Penetrating Delivery of Compounds and Nanoparticles into Tumors, Cancer Cell. 16 (2009) 510–520. https://doi.org/10.1016/j.ccr.2009.10.013.

[16] T. Teesalu, K.N. Sugahara, V.R. Kotamraju, E. Ruoslahti, C-end rule peptides mediate neuropilin-1-dependent cell, vascular, and tissue penetration, Proc. Natl. Acad. Sci. U. S. A. 106 (2009) 16157–16162. https://doi.org/10.1073/pnas.0908201106.

[17] K.N. Sugahara, P. Scodeller, G.B. Braun, T.H. De Mendoza, C.M. Yamazaki, M.D. Kluger, J. Kitayama, E. Alvarez, S.B. Howell, T. Teesalu, E. Ruoslahti, A.M. Lowy, A tumor-penetrating peptide enhances circulation-independent targeting of peritoneal carcinomatosis, J. Control. Release. 212 (2015) 59–69. https://doi.org/10.1016/j.jconrel.2015.06.009.

[18] J. Shi, P.W. Kantoff, R. Wooster, O.C. Farokhzad, Cancer nanomedicine: Progress, challenges and opportunities, Nat. Rev. Cancer. 17 (2017) 20–37. https://doi.org/10.1038/nrc.2016.108.

[19] I.M. Adjei, M.N. Temples, S.B. Brown, B. Sharma, Targeted nanomedicine to treat bone metastasis, Pharmaceutics. 10 (2018). https://doi.org/10.3390/pharmaceutics10040205.

[20] A. Yousefi, G. Storm, R. Schiffelers, E. Mastrobattista, Trends in polymeric delivery of nucleic acids to tumors, J. Control. Release. 170 (2013) 209–218. https://doi.org/10.1016/j.jconrel.2013.05.040.

[21] Guan-Hypoxia-induced tumor cell resistance-liposomes-Nanoscale2007.pdf, (n.d.).

[22] O. Paecharoenchai, N. Niyomtham, A. Apirakaramwong, T. Ngawhirunpat, T. Rojanarata, B.E. Yingyongnarongkul, P. Opanasopit, Structure relationship of cationic lipids on gene transfection mediated by cationic liposomes, AAPS PharmSciTech. 13 (2012) 1302–1308. https://doi.org/10.1208/s12249-012-9857-5.

[23] J.M. López-Pinto, M.L. González-Rodríguez, A.M. Rabasco, Effect of cholesterol and ethanol on dermal delivery from DPPC liposomes, Int. J. Pharm. 298 (2005) 1–12. https://doi.org/10.1016/j.ijpharm.2005.02.021.

[24] Y. Li, M. Alsagabi, D. Fan, G.S. Bova, A.H. Tewfik, S.M. Dehm, Intragenic rearrangement and altered RNA splicing of the androgen receptor in a cell-based model of prostate cancer progression, Cancer Res. 71 (2011) 2108–2117. https://doi.org/10.1158/0008-5472.CAN-10-1998.

[25] K.N. Sugahara, T. Teesalu, P. Prakash Karmali, V. Ramana Kotamraju, L. Agemy, D.R. Greenwald, E. Ruoslahti, Coadministration of a tumor-penetrating peptide enhances the efficacy of cancer drugs, Science (80-.). 328 (2010) 1031–1035. https://doi.org/10.1126/science.1183057.

[26] A. Van Bokhoven, M. Varella-Garcia, C. Korch, W.U. Johannes, E.E. Smith, H.L. Miller, S.K. Nordeen, G.J. Miller, M.S. Lucia, Molecular Characterization of Human Prostate Carcinoma Cell Lines, Prostate. 57 (2003) 205–225. https://doi.org/10.1002/pros.10290.

[27] J.K. Simmons, B.E. Hildreth, W. Supsavhad, S.M. Elshafae, B.B. Hassan, W.P. Dirksen, R.E. Toribio, T.J. Rosol, Animal Models of Bone Metastasis, Vet. Pathol. 52 (2015) 827–841. https://doi.org/10.1177/0300985815586223.

[28] M.E. Gleave, B.P. Monia, Antisense therapy for cancer, Nat. Rev. Cancer. 5 (2005) 468–479. https://doi.org/10.1038/nrc1631.

[29] S.D. Li, L. Huang, Targeted delivery of antisense oligodeoxynucleotide and small interference RNA into lung cancer cells, Mol. Pharm. 3 (2006) 579–588. https://doi.org/10.1021/mp060039w.

[30] S.K. Golombek, J.N. May, B. Theek, L. Appold, N. Drude, F. Kiessling, T. Lammers, Tumor targeting via EPR: Strategies to enhance patient responses, Adv. Drug Deliv. Rev. 130 (2018) 17–38. https://doi.org/10.1016/j.addr.2018.07.007.

[31] S. Sindhwani, A.M. Syed, J. Ngai, B.R. Kingston, L. Maiorino, J. Rothschild, P. MacMillan, Y. Zhang, N.U. Rajesh, T. Hoang, J.L.Y. Wu, S. Wilhelm, A. Zilman, S. Gadde, A. Sulaiman, B. Ouyang, Z. Lin, L. Wang, M. Egeblad, W.C.W. Chan, The entry of nanoparticles into solid tumours, Nat. Mater. 19 (2020) 566–575. https://doi.org/10.1038/s41563-019-0566-2.

[32] X. Liu, P. Lin, I. Perrett, J. Lin, Y.P. Liao, C.H. Chang, J. Jiang, N. Wu, T. Donahue, Z. Wainberg, A.E. Nel, H. Meng, Tumor-penetrating peptide enhances transcytosis of silicasome-based chemotherapy for pancreatic cancer, J. Clin. Invest. 127 (2017) 2007–2018. https://doi.org/10.1172/JCI92284.

[33] H.I. Scher, G. Buchanan, W. Gerald, L.M. Butler, W.D. Tilley, Targeting the androgen receptor: Improving outcomes for castration-resistant prostate cancer, Endocr. Relat. Cancer. 11 (2004) 459–476. https://doi.org/10.1677/erc.1.00525.

[34] I. Coutinho, T.K. Day, W.D. Tilley, L.A. Selth, Androgen receptor signaling in castration-resistant prostate cancer: A lesson in persistence, Endocr. Relat. Cancer. 23 (2016) T179–T197. https://doi.org/10.1530/ERC-16-0422.

[35] T. Visakorpi, E. Hyytinen, P. Koivisto, M. Tanner, R. Keinanen, C. Palmberg, A. Palotie, T. Tammela, J. Isola, O. Kallioniemi, Androgen Receptor Gene and Cancer, Nat. Genet. 9 (1995) 401–406.

[36] W.D. Tilley, G. Buchanan, T.E. Hickey, J.M. Bentel, Mutations in the androgen receptor gene are associated with progression of human prostate cancer to androgen independence, Clin. Cancer Res. 2 (1996) 277–285.

[37] C.J. Ryan, M.R. Smith, J.S. De Bono, A. Molina, C.J. Logothetis, P. De Souza, K. Fizazi, P. Mainwaring, J.M. Piulats, S. Ng, J. Carles, P.F.A. Mulders, E. Basch, E.J. Small, F. Saad, D. Schrijvers, H. Van Poppel, S.D. Mukherjee, H. Suttmann, W.R. Gerritsen, T.W. Flaig, D.J. George, E.Y. Yu, E. Efstathiou, A. Pantuck, E. Winquist, C.S. Higano, M.E. Taplin, Y. Park, T. Kheoh, T. Griffin, H.I. Scher, D.E. Rathkopf, Abiraterone in metastatic prostate cancer without previous chemotherapy, N. Engl. J. Med. 368 (2013) 138–148. https://doi.org/10.1056/NEJMoa1209096.

[38] T.M. Beer, A.J. Armstrong, D.E. Rathkopf, Y. Loriot, C.N. Sternberg, C.S. Higano, P. Iversen, S. Bhattacharya, J. Carles, S. Chowdhury, I.D. Davis, J.S. De Bono, C.P. Evans, K. Fizazi, A.M. Joshua, C.S. Kim, G. Kimura, P. Mainwaring, H. Mansbach, K. Miller, S.B. Noonberg, F. Perabo, D. Phung, F. Saad, H.I. Scher, M.E. Taplin, P.M. Venner, B. Tombal, Enzalutamide in metastatic prostate cancer before chemotherapy, N. Engl. J. Med. 371 (2014) 424–433. https://doi.org/10.1056/NEJMoa1405095.

[39] Y. Li, R. Yang, C.M. Henzler, Y. Ho, C. Passow, B. Auch, S. Carreira, D.N. Rodrigues, C. Bertan, T.H. Hwang, D.A. Quigley, H.X. Dang, C. Morrissey, M. Fraser, S.R. Plymate, C.A. PlymateMaher, F.Y. Feng, J.S. De Bono, S.M. Dehm, Diverse AR gene rearrangements mediate resistance to androgen receptor inhibitors in metastatic prostate cancer, Clin. Cancer Res. 26 (2020) 1965–1976. https://doi.org/10.1158/1078-0432.CCR-19-3023.

[40] D.A. Quigley, H.X. Dang, S.G. Zhao, P. Lloyd, R. Aggarwal, J.J. Alumkal, A. Foye, V. Kothari, M.D. Perry, A.M. Bailey, D. Playdle, T.J. Barnard, L. Zhang, J. Zhang, J.F. Youngren, M.P. Cieslik, A. Parolia, T.M. Beer, G. Thomas, K.N. Chi, M. Gleave, N.A. Lack, A. Zoubeidi, R.E. Reiter, M.B. Rettig, O. Witte, C.J. Ryan, L. Fong, W. Kim, T. Friedlander, J. Chou, H. Li, R. Das, H. Li, R. Moussavi-Baygi, H. Goodarzi, L.A. Gilbert, P.N. Lara, C.P. Evans, T.C. Goldstein, J.M. Stuart, S.A. Tomlins, D.E. Spratt, R.K. Cheetham, D.T. Cheng, K. Farh, J.S. Gehring, J. Hakenberg, A. Liao, P.G. Febbo, J. Shon, B. Sickler, S. Batzoglou, K.E. Knudsen, H.H. He, J. Huang, A.W. Wyatt, S.M. Dehm, A. Ashworth, A.M. Chinnaiyan, C.A. Maher, E.J. Small, F.Y. Feng, Genomic Hallmarks and Structural Variation in Metastatic Prostate Cancer, Cell. 174 (2018) 758–769.e9. https://doi.org/10.1016/j.cell.2018.06.039.

[41] K.T. Tietz, S.M. Dehm, Androgen receptor variants: RNA-based mechanisms and therapeutic targets, Hum. Mol. Genet. 29 (2020) R19–R26. https://doi.org/10.1093/hmg/ddaa089.

[42] N.-V. Mohamad, I.-N. Soelaiman, K.-Y. Chin, Clinical Interventions in Aging Dovepress A concise review of testosterone and bone health, Clin. Interv. Aging. (2016) 11–1317. https://doi.org/10.2147/CIA.S115472.

[43] M. Velders, P. Diel, How sex hormones promote skeletal muscle regeneration, Sport. Med. 43 (2013) 1089–1100. https://doi.org/10.1007/s40279-013-0081-6.

[44] K.T. Nead, G. Gaskin, C. Chester, S. Swisher-McClure, N.J. Leeper, N.H. Shah, Association between androgen deprivation therapy and risk of dementia, JAMA Oncol. 3 (2017) 49–55. https://doi.org/10.1001/jamaoncol.2016.3662.

[45] I.M. Adjei, B. Sharma, C. Peetla, V. Labhasetwar, Inhibition of bone loss with surface-modulated, drug-loaded nanoparticles in an intraosseous model of prostate cancer, J. Control. Release. 232 (2016) 83–92. https://doi.org/10.1016/j.jconrel.2016.04.019.

[46] M. Kawatani, H. Osada, Osteoclast-targeting small molecules for the treatment of neoplastic bone metastases, Cancer Sci. 100 (2009) 1999–2005. https://doi.org/10.1111/j.1349-7006.2009.01294.x.

[47] M. Esposito, Y. Kang, Targeting tumor-stromal interactions in bone metastasis, Pharmacol. Ther. 141 (2014) 222–233. https://doi.org/10.1016/j.pharmthera.2013.10.006.

[48] S.I. Thamake, S.L. Raut, Z. Gryczynski, A.P. Ranjan, J.K. Vishwanatha, Alendronate coated poly-lactic-co-glycolic acid (PLGA) nanoparticles for active targeting of metastatic breast cancer, Biomaterials. 33 (2012) 7164–7173. https://doi.org/10.1016/j.biomaterials.2012.06.026.

[49] L.E. Cole, T. Vargo-Gogola, R.K. Roeder, Targeted delivery to bone and mineral deposits using bisphosphonate ligands, Adv. Drug Deliv. Rev. 99 (2016) 12–27. https://doi.org/10.1016/j.addr.2015.10.005.

[50] M. Neudert, C. Fischer, B. Krempien, F. Bauss, M.J. Seibel, Site-specific human breast cancer (MDA-MB-231) metastases in nude rats: Model characterisation and in vivo effects of ibandronate on tumour growth, Int. J. Cancer. 107 (2003) 468–477. https://doi.org/10.1002/ijc.11397.

[51] E. Corey, L.G. Brown, J.E. Quinn, M. Poot, M.P. Roudier, C.S. Higano, R.L. Vessella, Zoledronic acid exhibits inhibitory effects on osteoblastic and osteolytic metastases of prostate cancer, Clin. Cancer Res. 9 (2003) 295–306.

[52] K. Ramanlal Chaudhari, A. Kumar, V.K. Megraj Khandelwal, M. Ukawala, A.S. Manjappa, A.K. Mishra, J. Monkkonen, R.S. Ramachandra Murthy, Bone metastasis targeting: A novel approach to reach bone using Zoledronate anchored PLGA nanoparticle as carrier system loaded with Docetaxel, J. Control. Release. 158 (2012) 470–478. https://doi.org/10.1016/j.jconrel.2011.11.020.

[53] K.B. Farrell, A. Karpeisky, D.H. Thamm, S. Zinnen, Bisphosphonate conjugation for bone specific drug targeting, Bone Reports. 9 (2018) 47–60. https://doi.org/10.1016/j.bonr.2018.06.007.

[54] E.J. Carbone, K. Rajpura, B.N. Allen, E. Cheng, B.D. Ulery, K.W.H. Lo, Osteotropic nanoscale drug delivery systems based on small molecule bone-targeting moieties, Nanomedicine Nanotechnology, Biol. Med. 13 (2017) 37–47. https://doi.org/10.1016/j.nano.2016.08.015.

[55] F. Wang, L. Chen, R. Zhang, Z. Chen, L. Zhu, RGD peptide conjugated liposomal drug delivery system for enhance therapeutic efficacy in treating bone metastasis from prostate cancer, J. Control. Release. 196 (2014) 222–233. https://doi.org/10.1016/j.jconrel.2014.10.012.

