## Supplemental Data for "iRGD-liposomes enhance tumor delivery and therapeutic efficacy of antisense oligonucleotide drugs against primary prostate cancer and bone metastasis"

**Supplementary Table S1** The physicochemical properties of the nanoparticles.

| Physicochemical properties | iRGD-liposome-ASO 1 | iRGD-liposome-ASO 2 | Liposome-ASO |
|---|---|---|---|
| Mean Sizes (nm) | 150 ± 36 | 210 ± 83 | 141 ± 36 |
| PDI | 0.0838 | 0.0850 | 0.0863 |
| Zeta (mV) | -6.67 ± 0.18 | -7.10 ± 0.42 | -14.4 ± 7.25 |
| ASO Conc. (mg/mL) | 8.093 ± 0.674 | 7.434 ± 0.804 | 7.368 ± 0.951 |

Data are presented as the mean ± standard deviation (n=3).

Abbreviations: ASO, antisense oligonucleotide; PDI, polydispersity index; conc., concentration.

**Supplementary Table S2** Pharmacokinetic parameters of ASO after intravenous administration of iRGD-liposome-ASO, liposome-ASO and ASO to mice (50 mg/kg per mouse).

| **Pharmacokinetic parameters** | **iRGD-liposome-ASO** | **Liposome-ASO** | **ASO** |
|---|---|---|---|
| $T_{1/2}$ (h) | 0.937 ± 0.014 | 0.918 ± 0.040 | 0.544 ± 0.004 |
| $AUC_{0\text{-}t}$ | 99641.44 ± 6815.08 * | 98834.77 ± 5765.90 * | 34474.02 ± 2062.38 |
| $F_r$ (%) | 289.03 ± 19.77 | 286.69 ± 16.73 | - |

Values are expressed as mean ± SD (n=3), * P<0.05 relative to ASO

Abbreviations: $T_{1/2}$: half-life; $AUC_{0\text{-}t}$: area under the concentration-time curve; $F_r$: relative bioavailability

Fig. S1

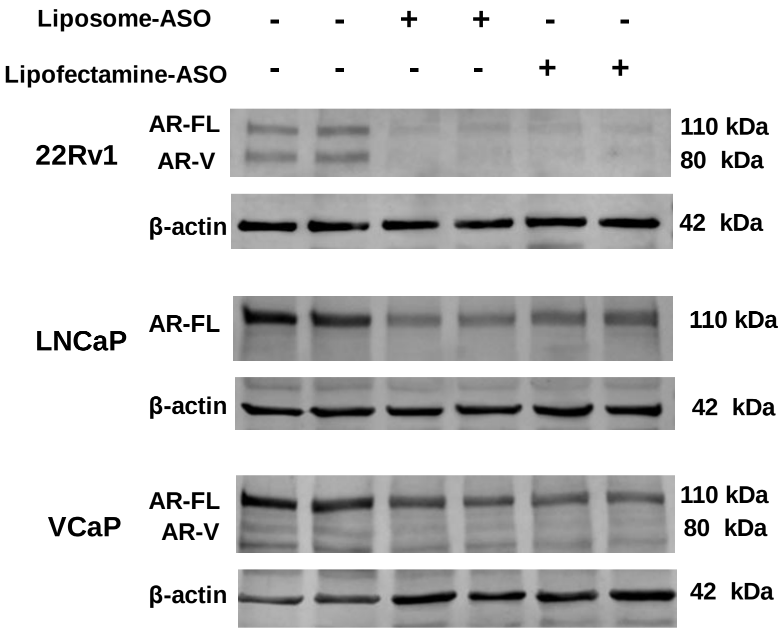


**Fig. S1. Inhibition of AR by liposome-ASO in vitro.**

The *in vitro* knockdown of AR-FL in 22Rv1, LNCaP, and VCaP cell lines at the protein levels determined by Western Blotting with the incubation of liposome-ASO or lipofectamine-ASO. The same amount of ASO (5 μg for 1 x $10^6$ cells) in the lipofectamine and the liposome formulations was used in each experiment. AR-FL: full-length androgen receptor; AR-V: AR splice variant.

Fig. S2

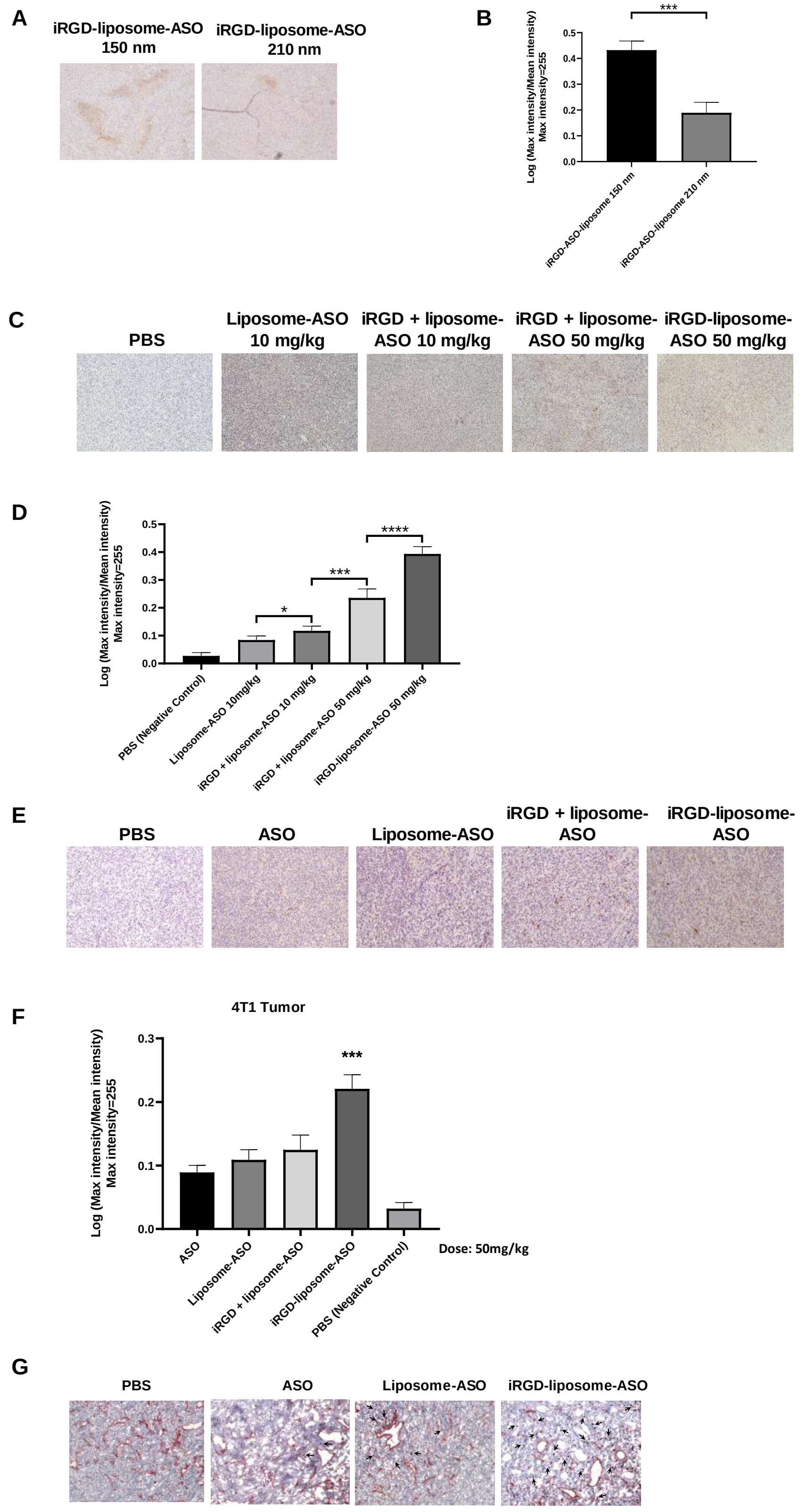
A
iRGD-liposome-ASO 150 nm
iRGD-liposome-ASO 210 nm
B
***
Log (Max intensity/Mean intensity)
Max intensity=255
0.0
0.1
0.2
0.3
0.4
0.5
iRGD-ASO-liposome 150 nm
iRGD-ASO-liposome 210 nm
C
PBS
Liposome-ASO 10 mg/kg
iRGD + liposome-ASO 10 mg/kg
iRGD + liposome-ASO 50 mg/kg
iRGD-liposome-ASO 50 mg/kg
D
Log (Max intensity/Mean intensity)
Max intensity=255
0.0
0.1
0.2
0.3
0.4
0.5
*
***
****
PBS (Negative Control)
Liposome-ASO 10mg/kg
iRGD + liposome-ASO 10 mg/kg
iRGD + liposome-ASO 50 mg/kg
iRGD-liposome-ASO 50 mg/kg
E
PBS
ASO
Liposome-ASO
iRGD + liposome-ASO
iRGD-liposome-ASO
F
4T1 Tumor
Log (Max intensity/Mean intensity)
Max intensity=255
0.0
0.1
0.2
0.3
***
ASO
Liposome-ASO
iRGD + liposome-ASO
iRGD-liposome-ASO
PBS (Negative Control)
Dose: 50mg/kg
G
PBS
ASO
Liposome-ASO
iRGD-liposome-ASO

**Fig. S2. iRGD-liposome enhances homing and uptake of ASO in 4T1 tumor model.**

(A) Representative immunohistochemistry (IHC) images and (B) quantification of ASO staining in tumors from mice bearing 4T1 orthotopic breast tumors. The tumors were collected after 2 h in vivo homing by different sizes of iRGD-liposomes with 10 mg/kg dosage of ASO in mice bearing 4T1 breast tumors. (C) Representative IHC images and (D) quantification of ASO staining in tumors from mice bearing 4T1 orthotopic breast tumors. The tumors were collected 4 h after different doses of ASO (10 mg/kg and 50 mg/kg) using iRGD covalently conjugated liposomes (iRGD-liposome-ASO) and unconjugated iRGD and liposomes (iRGD + liposome-ASO), respectively. (E) Representative IHC sections and (F) quantification of ASO staining in 4T1 orthotopic breast tumors. The tumors were collected after 4-day homing by different delivery systems including PBS, free ASO, liposome-ASO, iRGD + liposome-ASO, and iRGD-liposome-ASO. The ASO dose was 50mg/kg. The density of ASO expression were analyzed by ImageJ. (G) IHC images of ASO accumulation in tumor stained with anti-ASO antibody for ASO (arrows) and anti-CD31 for blood vessels (red) after 2 h in vivo homing. The ASO dose was 50mg/kg. All experiments were performed in three mice per group. The data are represented as mean ± standard deviation (SD). *P<0.05, ***P<0.001, ****P<0.0001 vs ASO.

Fig. S3

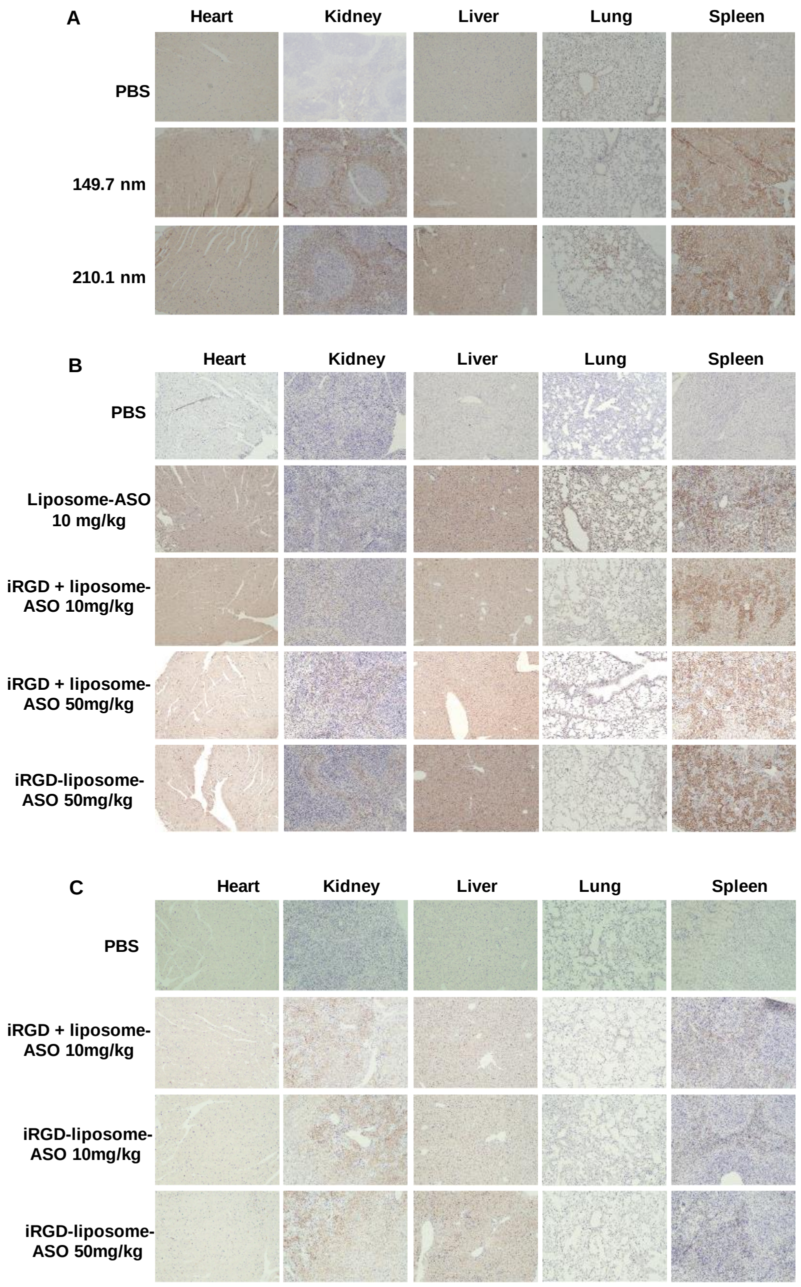


**Fig. S3. Distribution of iRGD-liposome-ASO in 4T1 tumor model.**

(A) IHC images of ASO staining in tissues collected after 2 h in vivo homing by different sizes of iRGD-liposomes with 10 mg/kg dosage of ASO. (B) IHC images of ASO staining in tissues collected 4 h after different doses of ASO (10 mg/kg and 50 mg/kg) using iRGD covalently conjugated liposomes (iRGD-liposome-ASO) and unconjugated iRGD and liposomes (iRGD + liposome-ASO), respectively. (C) ASO staining in tissues after 4-day homing by different delivery systems. All experiments were performed in three mice per group; representative images are shown here.

Fig. S4

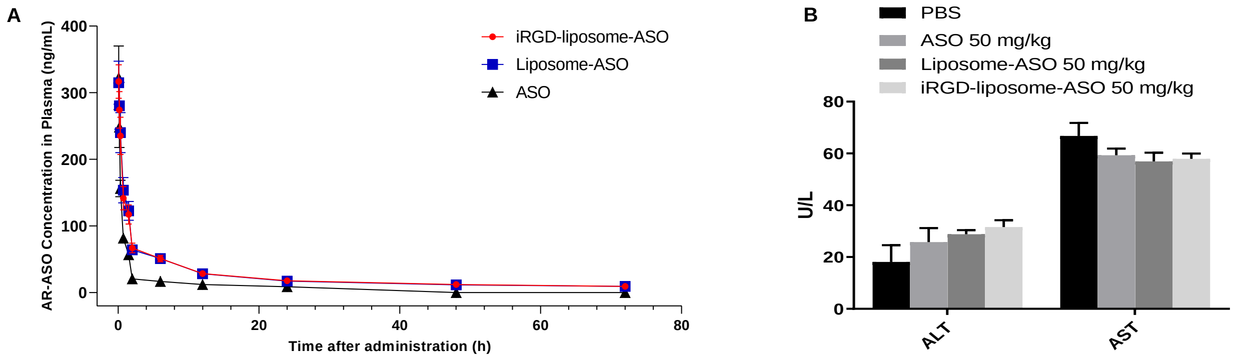


**Fig. S4. Pharmacokinetics and biological safety of iRGD-liposome-ASO *in vivo*.**

(A) Mean plasma concentration-time curves of AR-ASO following intravenous administration of ASO, liposome-ASO and iRGD-liposome-ASO. (B) Plasma levels of liver enzymes aspartate transaminase (AST) and alanine transaminase (ALT) in 22Rv1-bearing mice. The mice were treated with a 50mg/kg dosage of ASO by different delivery system every 4 days for two weeks. The data are presented as the mean ± standard deviation (SD). Statistical significance was calculated using Student’s t test.

Fig. S5

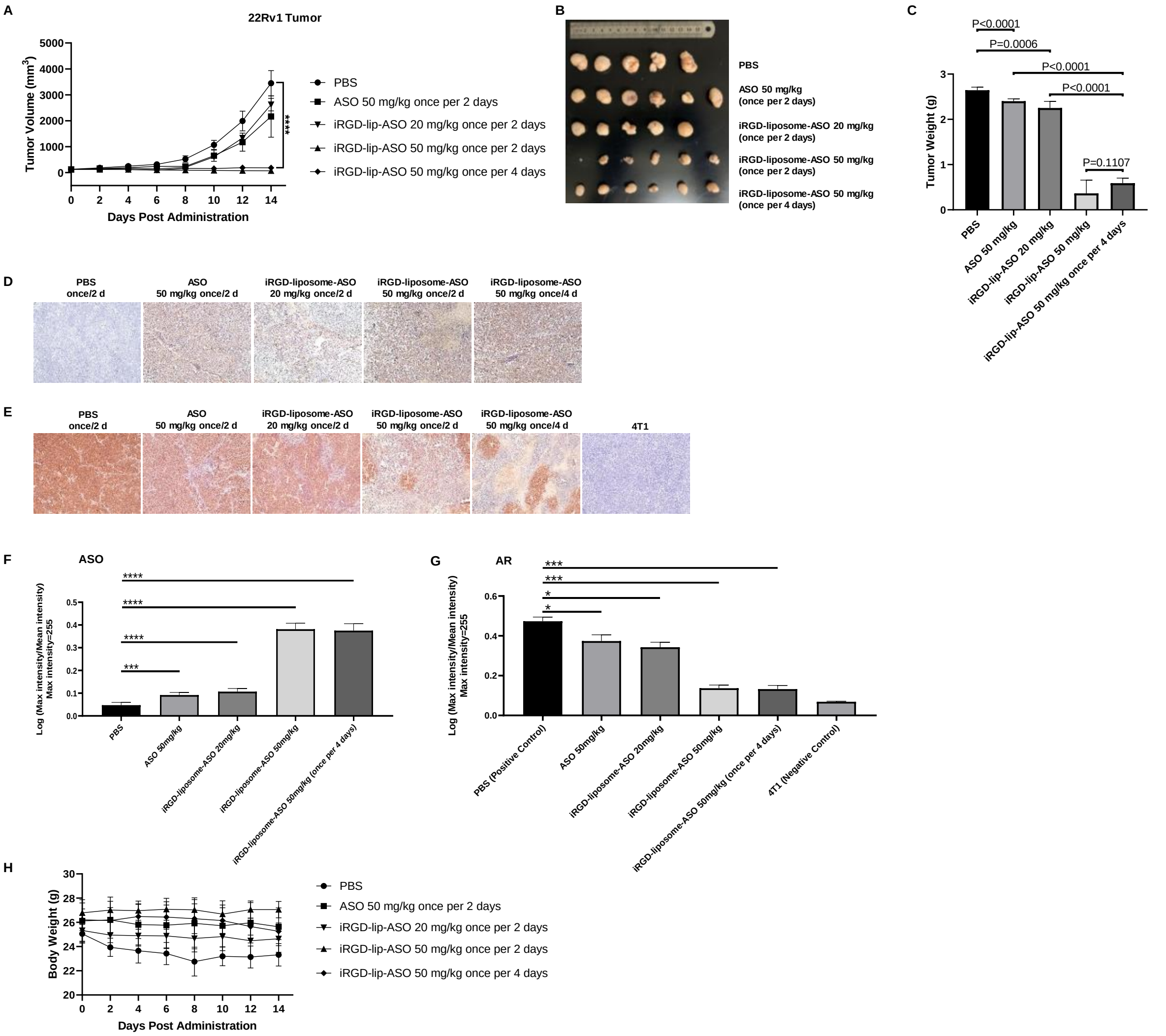


**Fig. S5. In vivo pharmacodynamics for therapeutic efficacy of iRGD-liposome-ASO in CRPC 22Rv1 tumor.**

The 22Rv1 tumor-bearing mice were treated with PBS, free ASO, liposome-ASO, and iRGD-liposome-ASO, respectively. The ASO dose was 20 mg/kg or 50 mg/kg and the injections were given every 2 or 4 days for two weeks, respectively. (A) Tumor volume growth curves of 22Rv1 tumor-bearing mice treated with different treatments for two weeks. (B) Images of tumor and (C) average tumor weight of mice bearing 22Rv1 tumor at the end of treatment. (D) Representative images of ASO staining by IHC in tumors after different treatments. (E) Representative IHC images of AR staining in tumors after different treatments. AR staining in 4T1 tumors serves as the negative control. (F) Quantification of ASO density in IHC sections from tumors with different treatments. (G) Quantification of AR density in IHC sections from tumors with different treatments. (H) Body weight in mice receiving different treatments over time. The density of ASO and AR expression were analyzed by ImageJ. All experiments were performed in 5-6 mice per group. The data are represented as mean ± standard deviation (SD). *P<0.05, ***P<0.001, **** P<0.0001 vs PBS controls.

Fig. S6

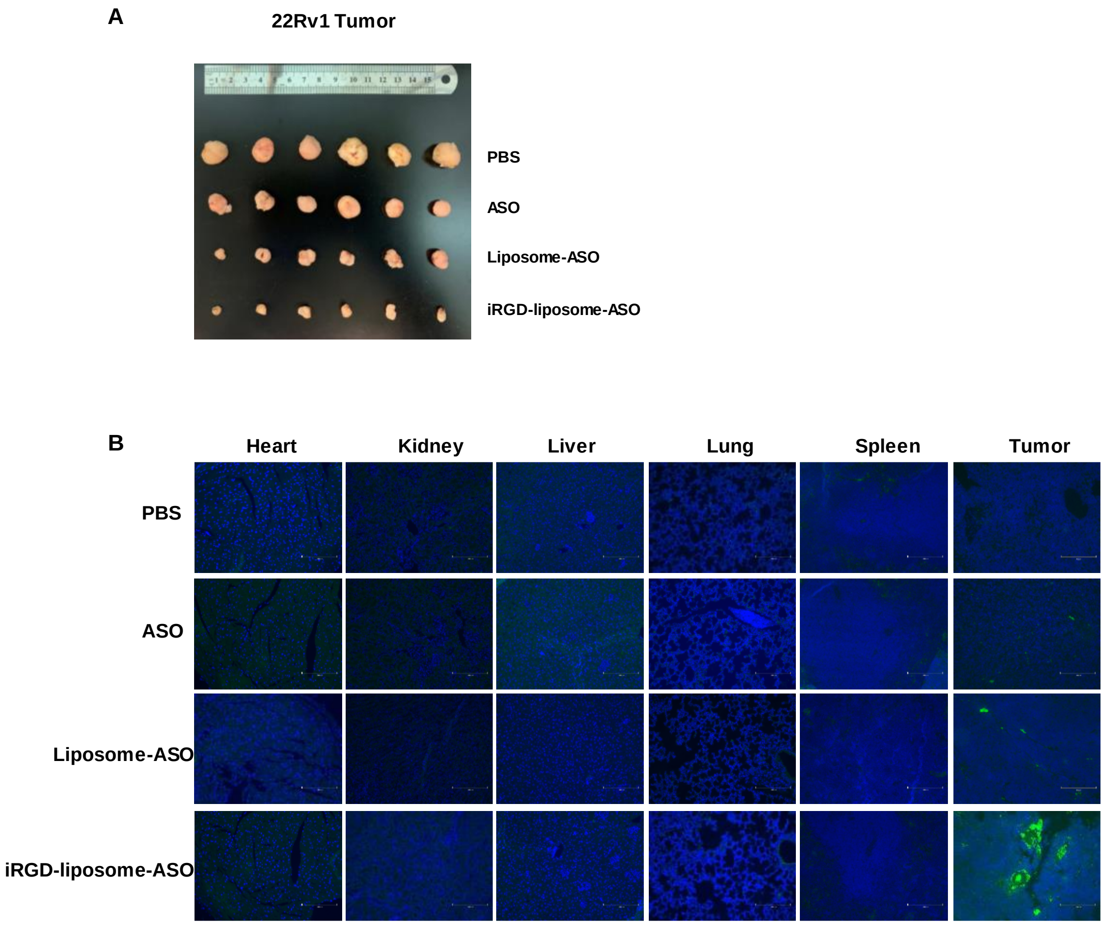


**Fig. S6. Short-term therapeutic efficacy of iRGD-liposome-ASO in CRPC 22Rv1 tumor.**

The 22Rv1 tumor-bearing mice were treated with PBS, free ASO, liposome-ASO, and iRGD-liposome-ASO, respectively. The ASO dose was 50 mg/kg and the injections were given every 4 days for two weeks, respectively. (A) Images of tumor of mice bearing 22Rv1 subcutaneous xenografts at the end of treatment. (B) IF images of TUNEL assay showing apoptotic cells in tumor and other tissues of 22Rv1-bearing mice with a 50mg/kg dosage of ASO every 4 days for two weeks. All experiments were performed in 5-6 mice per group.

Fig. S7

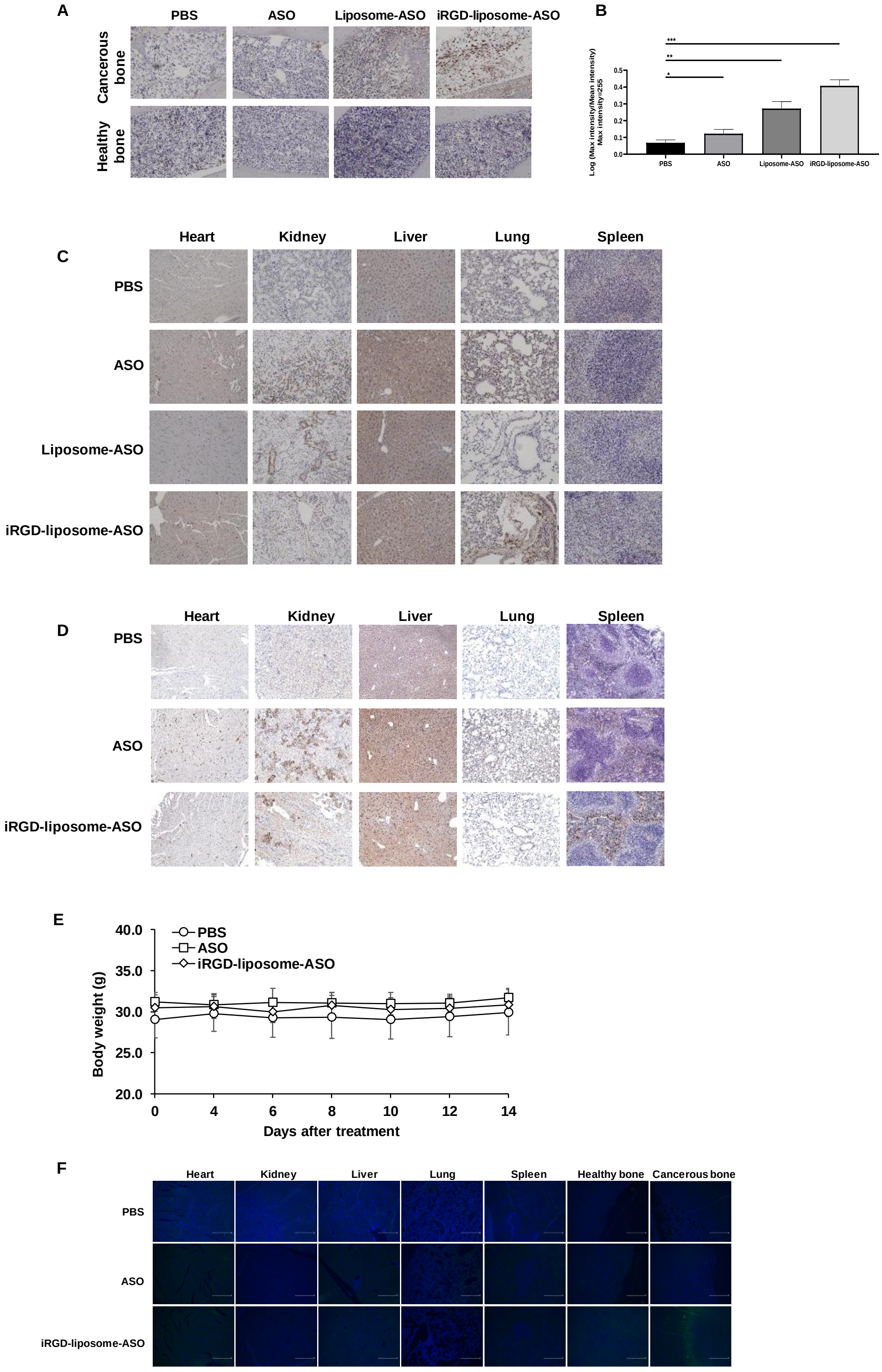
A
PBS
ASO
Liposome-ASO
iRGD-liposome-ASO
Cancerous bone
Healthy bone
B
Log (Max intensity/Mean intensity) Max intensity=255
PBS
ASO
Liposome-ASO
iRGD-liposome-ASO
C
Heart
Kidney
Liver
Lung
Spleen
PBS
ASO
Liposome-ASO
iRGD-liposome-ASO
D
Heart
Kidney
Liver
Lung
Spleen
PBS
ASO
iRGD-liposome-ASO
E
PBS
ASO
iRGD-liposome-ASO
Body weight (g)
Days after treatment
F
Heart
Kidney
Liver
Lung
Spleen
Healthy bone
Cancerous bone
PBS
ASO
iRGD-liposome-ASO

**Fig. S7. iRGD-liposome-ASO homing and treatment in the CRPC bone metastasis model.**

(A) IHC sections of ASO staining and (B) quantification of ASO density in IHC sections of bones from healthy mice and those with bone metastasis. Bones were collected 2 h after injection with PBS, ASO, liposome-ASO and iRGD-liposome-ASO, respectively. The ASO dose was 50 mg/kg. (C) Representative IHC sections of ASO staining in tissues after 2-hour homing by different treatments. The ASO dose was 50 mg/kg. (D) Representative IHC sections of ASO staining in tissues by the end of the treatments. The mice bearing bone metastasis were treated with PBS, free ASO, and iRGD-liposome-ASO, respectively. The ASO dose was 50mg/kg and the injections were given every 2 days for two weeks. (E) Body weight in mice receiving different treatments over time. (F) IF images of TUNEL assay showing apoptotic cells in healthy and cancerous bone as well as other tissues with different treatments. The ASO dose was 50mg/kg and the injections were given every 2 days for two weeks. The density of ASO expression were analyzed by ImageJ. The data are represented as mean ± standard deviation (SD). *$P<0.05$, **$P<0.01$, ***$P<0.001$ vs PBS controls.
